# Disease spectrum and outcomes among elderly patients in two tertiary hospitals in Dar es Salaam, Tanzania

**DOI:** 10.1101/554972

**Authors:** Basil Tumaini, Patricia Munseri, Kisali Pallangyo

## Abstract

**Background:** There has been an increase in the number of individuals aged ≥60 years in Tanzania and in sub Saharan Africa in general due to improved survival. However there is scarcity of data on the disease burden, patterns and outcomes following admission in this population. We therefore describe the pattern of diagnoses, outcomes and factors associated with the outcomes among elderly patients admitted at Muhimbili National Hospital (MNH) and Jakaya Kikwete Cardiac Institute (JKCI) medical wards.

**Methodology:** We prospectively enrolled patients aged ≥60 years (elderly) admitted through MNH Emergency Medicine to the MNH medical wards and JKCI. ICD 10 was used to code for disease diagnosis at discharge or death. Modified Barthel index was used to assess for functional activity on admission and at discharge.

**Results:** We enrolled 336 elderly participants, who comprised 30.1% of all medical admissions. The mean age was 70.6 years; 50% were female and 263 (78.3%) had comorbidities with an average of 2 diagnoses per participant. The most common diagnoses were: hypertension (44.9%), stroke (31.5%), heart failure (18.5%), pneumonia (17.9%), diabetes mellitus (17.3%) and chronic kidney disease (16.4%). The median duration of hospital stay was 5 days and in-hospital mortality was 25.6%. Non-communicable diseases (NCDs) accounted for 65% of the deaths and 50% of the deaths occurred within 72 hours of hospitalization. A modified Barthel score of ≤20 on admission was associated with 15 times increased risk of death (p<0.001).

**Conclusion:** NCDs were the most common diagnoses and cause of death among patients aged ≥60 years admitted to the medical wards. In-hospital mortality was high and occurred within 72 hours of hospitalization. A modified Barthel score <20 on admission was associated with mortality.

## Introduction

The average life expectancy of a Tanzanian at birth has improved from 49.6 years during the period of 1990 – 1995 to 66.7 years during the period of 2015 – 2020 largely due to success in controlling HIV/AIDS [1]. This, together with declining fertility, is resulting in population aging which is a worldwide phenomenon. Population aging is projected to have profound socioeconomic consequences in the not-too-distant future.

The proportion of elderly patients among medical admissions in developing countries has been shown to be increasing, [2–5]. Aging is a major risk factor underlying disease and disability; again, older people have altered responses to therapies developed for younger adults often with less effectiveness and more adverse reactions. The rapid increase in the number of older patients is predicted to overwhelm health care systems with their mostly chronic diseases and conditions [6]. Because of their complex medical problems, management of elderly patients is best done by a multidisciplinary team with high levels of personnel and requires adequate financial resources; input from geriatricians - who are scarce in many developing countries - is also required. Hence aging and old age poses significant challenges to medicine. Despite this being a highly dynamic group, it is still understudied in many developing countries and hence evidence for guiding policy interventions is lacking.

The pattern of diseases among geriatric patients admitted to the medical wards in developing countries has been evolving from predominantly infectious diseases to predominantly noncommunicable diseases [2,4,5,7]. This makes the periodic evaluation of the changing patterns imperative to guide practice and appropriate policies. It has also been shown that physical disability as well as impaired capacity for self-care increased rapidly with advancing age similar to comorbidity [2]. Hence evaluation of the functional status is an important element of any geriatric assessment. Evaluation of functional status using simple yet invaluable ADL tools is not part of the standard of care in Tanzania and hence the usability of these tools as well as information on the magnitude of disability is lacking.

The study aimed at investigating the contribution of individuals aged ≥60 years to the medical admissions; their demographics, diseases, hospital outcomes as well as associated factors, at two tertiary healthcare facilities in Dar es Salaam Tanzania and to compare such data with a study done about three decades ago in the same center. This has been achieved.

## Materials and methods

### Ethics statement

Ethical approval was obtained from the Muhimbili University of Health and Allied Sciences Institutional Review Board. Muhimbili National Hospital and Jakaya Kikwete Cardiac Institute administration granted permission to conduct the study in the respective institutions. Written informed consent was obtained from all study participants /caretakers for patients who were unable to provide consent prior to enrolment into the study. All participants received treatment as per hospital standard operating guidelines.

### Study design and population

This study was conducted at two public tertiary hospitals located in Dar es Salaam, Tanzania: Muhimbili National Hospital (MNH) and Jakaya Kikwete Cardiac Institute (JKCI). These hospitals have high skilled staff and advanced diagnostic facilities. We prospectively enrolled patients aged ≥60 years who were admitted by MNH Emergency Medical Department (EMD) to the medical wards at MNH and JKCI between October and December 2017.

### Data collection

The investigator (B.T.) administered interviewer-based structured questions to the study participant/caretaker for participants who were unable to communicate. Information on socio-demographic characteristics was collected and adaptation of the modified Barthel Index [8,9] was used to establish independence in the performance of the activities of daily living (ADL) at admission and discharge. Guidelines for the Barthel Index were employed to ensure what was recorded is what the patient does, using the best available evidence. Unconscious patients scored 0 throughout. Patients were followed up until the outcome: discharge from the hospital or death. Final diagnoses for each participant were recorded from the hospital charts and thereafter coded using ICD-10 [10] at the time of outcome. Length of hospital stay was computed as the difference between the date of admission and the date of discharge/death for each participant. The underlying cause of death was obtained from the death certificates. We studied factors such as socio-demographic characteristics, diagnoses and functional status on admission as measured by the modified Barthel ADL index and their association with mortality.

### Statistical methods

Data was entered into EpiData version 3.1 and thereafter exported into IBM® SPSS®Statistics version 23 for analysis. Qualitative variables such as sex, level of education, marital status, hospital outcome, discharge diagnosis, cause of death and health insurance status were summarized as frequency and proportions. Differences in proportions across groups were compared using Chi-square test or Fisher’s exact test. Quantitative variables such as length of hospital stay, age, and modified Barthel ADL score at admission and at discharge were summarized as means and standard deviation. The student’s t-test was used to compare quantitative variables among participants who survived to those who died. A logistic regression analysis was performed so as to identify independent risk factors for mortality. All factors with p value of <0.2 in univariate analysis were included in the multivariate analysis model. Statistical significance was accepted at p<0.05.

## Results

During the three-month study period, 1301 patients were admitted to the MNH and JKCI medical wards out of which 392 (30.1%) were aged ≥60 years. Details of enrolment are shown in Fig 1, in brief 56 (14%) patients were excluded; (13 patients were deemed to have surgical conditions and were transferred to surgical wards, 27 were unable to communicate and had no next of keen to provide informed consent and 16 patients refused to take part in the study). Therefore, a total of 336 participants were enrolled into the study.

**Fig 1.**
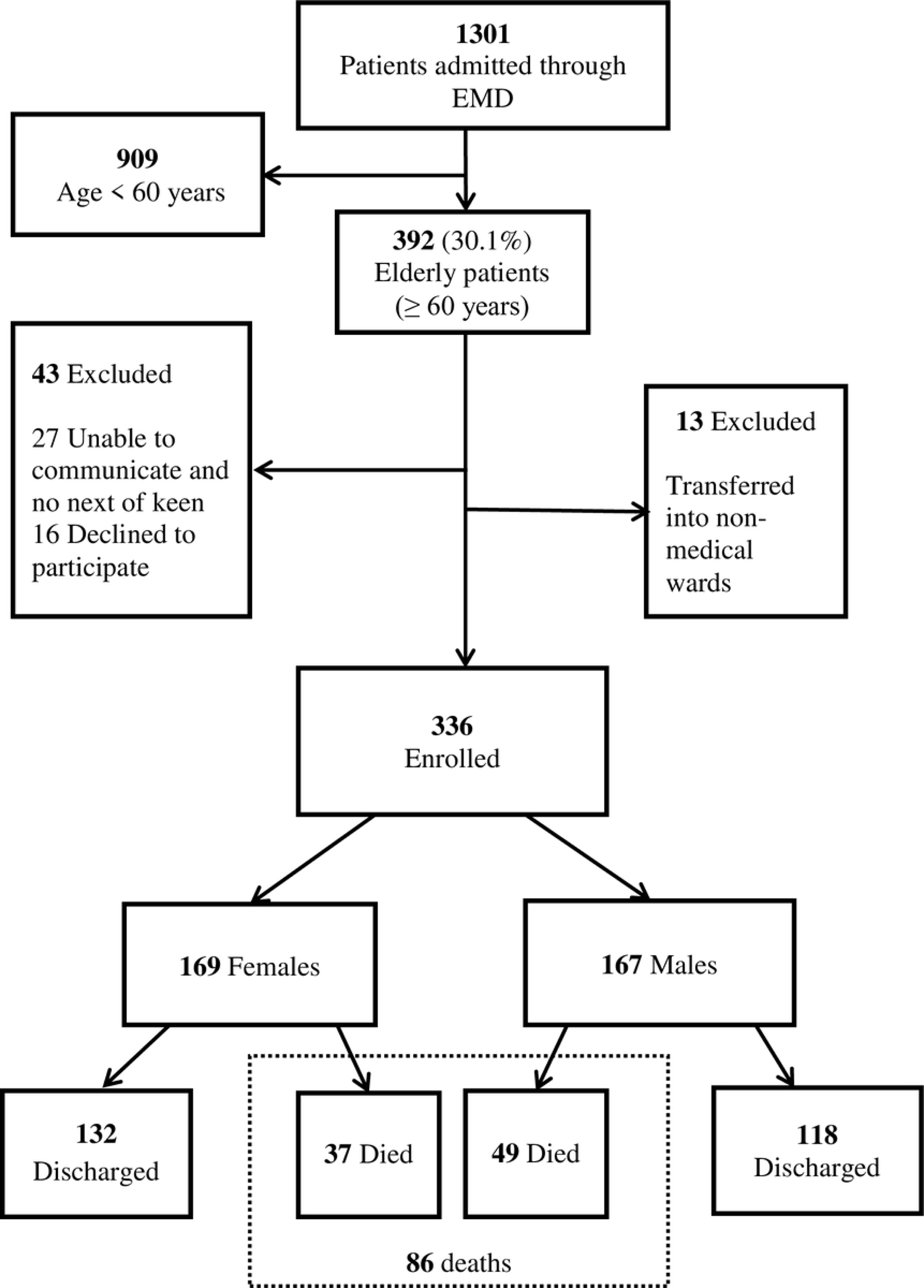
Consort diagram. EMD = Emergency medical department

### Socio-demographic characteristics of study participants

Table 1 summarizes the socio-demographic characteristics of the study participants. There were 169 (50.3%) females enrolled, the mean age (± SD) for enrolled participants was 70.6 (± 8.9) years, 180 (53.6%) were in the age group 60 – 69 years and 132 (39.3%) had health insurance. About half of the study participants had at least primary education however, there was a significant difference in proportion of uneducated females compared to males 45 (26.6%) vs. 19 (11.4%) respectively p<0.001.

**Table 1.**
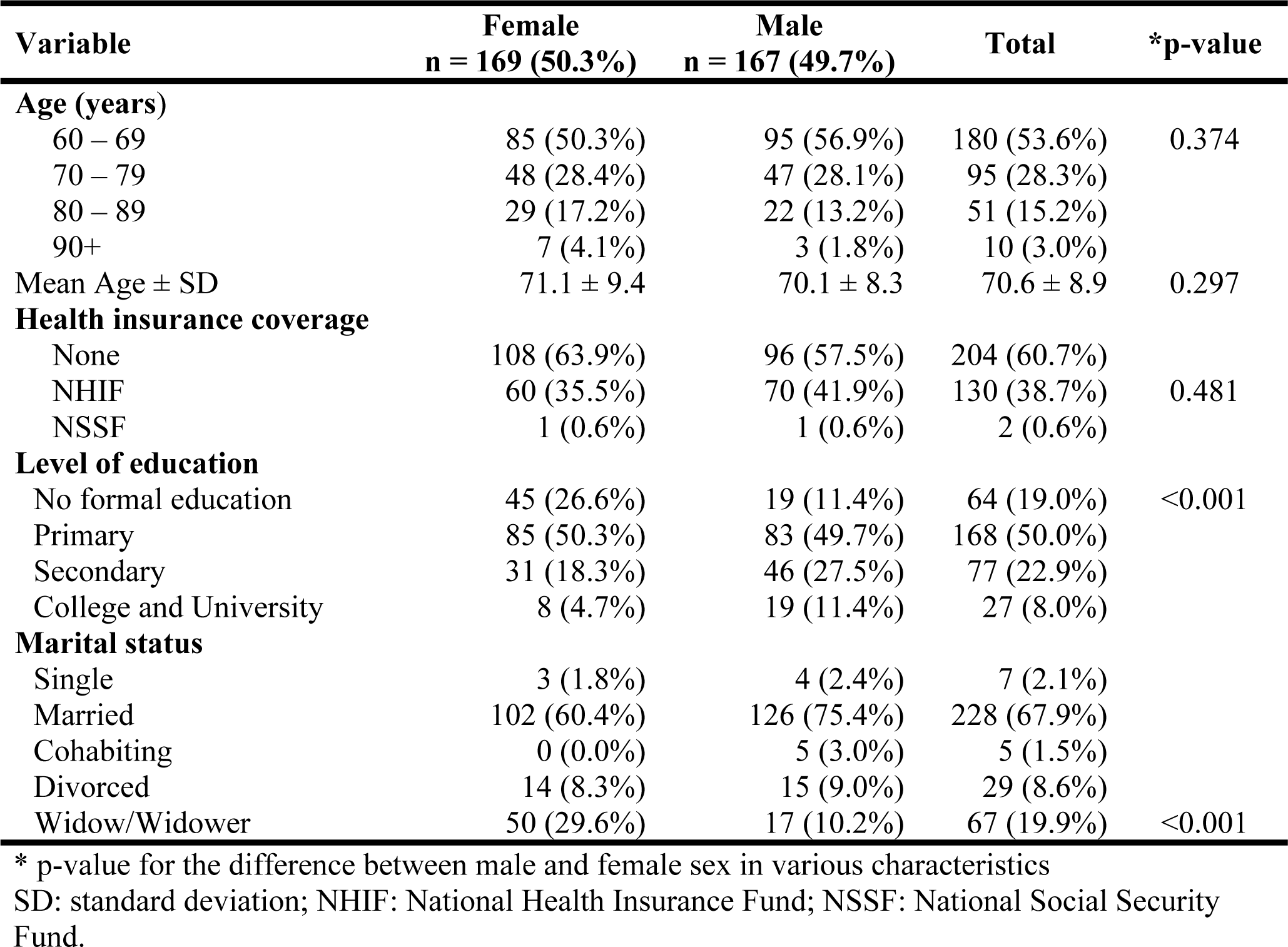
Socio-demographic characteristics of the elderly at admission (N=336).

### Diagnosis of the study participants

The description of diagnosis of the study participants are summarized in Table 2, 73 (21.7%) had single diagnosis while 263 (78.3%) had more than one diagnosis with an average of 2.3 diagnoses per study participant. The common diagnoses were hypertension 151 (44.9%), stroke 106 (31.5%), heart failure 62 (18.5%), pneumonia, 60 (17.9%), diabetes mellitus 58 (17.3%) and chronic kidney disease (CKD) 55 (16.4%). HIV was diagnosed in 4.8% (16/336) of the study participants. Frequent comorbid conditions were stroke and hypertension 73 (21.7%), diabetes mellitus and hypertension 38 (11.3%), CKD and diabetes mellitus or hypertension 34 (10.1%), stroke and pneumonia 32 (9.5%), and heart failure and hypertension in 28 participants (8.3%). Overall, the proportion of non-communicable diseases was 82.7%.

**Table 2.** Diagnoses of the study participants (N=336).

| S/N | ICD 10 codes** | Diagnoses | Cases (% of cases)* | Proportion of diagnoses |
| --- | --- | --- | --- | --- |
| 1 | I10 & I15 | Hypertension | 151 (44.9%) | 19.4% |
| 2 | I63 & I61 | Stroke (Ischemic, hemorrhagic) | 106 (31.5%) | 13.6% |
| 3 | I50.9, I25.5, I42.0 | Heart failure | 62 (18.5%) | 7.9% |
| 4 | J18 | Pneumonia | 60 (17.9%) | 7.7% |
| 5 | E11 | Type 2 diabetes mellitus | 58 (17.3%) | 7.4% |
| 6 | N18 | Chronic kidney disease (CKD) | 55 (16.4%) | 7.1% |
| 7 | C22 - 95.9 | Malignant neoplasms | 38 (11.3%) | 4.9% |
| 8 | D64.9 | Anemia | 36 (10.7%) | 4.6% |
| 9 | E87 | Electrolyte disorders | 20 (6.0%) | 2.6% |
| 10 | B20 | HIV disease | 16 (4.8%) | 2.1% |
| 11 | K27 | Peptic ulcer disease | 14 (4.2%) | 1.8% |
| 12 | E39.0 | Urinary tract infection | 11 (3.3%) | 1.4% |
| 13 | B54 | Malaria | 7 (2.1%) | 0.9% |
| 14 | A15-19 | Tuberculosis | 6 (1.8%) | 0.8% |
| 15 | B90 | Sequelae of tuberculosis | 5 (1.5%) | 0.6% |
|  | Multiple | Other diagnoses | 135 (40.2%) | 17.3% |
|  |  |  | 780 (232.10%) | 100% |

\* Some participants had more than one disease and hence the percentage of cases adds to more than 100%.
\*\* Refer Supporting Information (S1 Table) for ICD-10 codes employed to group the data.
HIV: human immunodeficiency virus disease.

### Hospital outcomes, duration of hospital stay and time to death

Of the 336 study participants, 250 (74.4%) were discharged home; 6 (1.8%) of these against medical advice. The overall in-hospital mortality was 86 (25.6%). The commonest underlying causes of mortality are shown in Table 3. Noncommunicable diseases were the cause of 56 (65%) of the deaths.

**Table 3.** Underlying cause of death by sex (N = 86).

| Underlying cause of death | Female | Male | Total | p-value |
| --- | --- | --- | --- | --- |
| Stroke | 9 (24.3%) | 16 (32.7%) | 25 (29.1%) | 0.400 |
| Pneumonia | 11 (29.7%) | 7 (14.3%) | 18 (20.9%) | 0.081 |
| Heart Failure | 10 (27.0%) | 8 (16.3%) | 18 (20.9%) | 0.227 |
| Chronic kidney disease | 2 (5.4%) | 5 (10.2%) | 7 (8.1%) | 0.694 |
| Sepsis* | 1 (2.7%) | 4 (8.2%) | 5 (5.8%) | 0.385 |
| Metabolic disorders | 1 (2.7%) | 3 (6.1%) | 4 (4.7%) | 0.631 |
| Meningitis | 2 (5.4%) | 1 (2.0%) | 3 (3.5%) | 0.575 |
| Malignant neoplasms | 0 (0.0%) | 2 (4.1%) | 2 (2.3%) | 0.504 |
| Other causes | 1 (2.7%) | 3 (6.1%) | 4 (4.7%) | 0.631 |
| **Total deaths** | **37 (21.9%)** | **49 (29.3%)** | **86 (25.6%)** | **0.118** |

\* Excluding pneumonia and meningitis

The median duration of hospital stay for the survivors was 5 days (range: 1 - 49 days), 162 (64.8%) participants were discharged within 7 days from admission and 222 (88.8%) were discharged within 14 days. Four study participants (1.6%) had a hospital stay of more than 4 weeks. The median time from admission to death was 3 days (range: 0 – 20 days), 48 (55.8%) deaths occurred in the first three days of hospitalization. Overall, 113 (33.6%) had a modified Barthel index of ≤20 (total dependence on performance of activities of daily living [ADL]) on admission. Of the 250 survivors, 48 (19.2%) had modified Barthel index of ≤20 at admission whereas 21 of these study participants (8.4%) were still totally dependent at discharge. The median modified Barthel score on admission among those who died was 12 (total dependence) whereas it was 63 (moderate dependence) among survivors, p< 0.001.

### Factors associated with the hospital outcomes

The mean age of study participants who died in hospital was 72.7 (±9.1) years compared to 69.9 (±8.7) years for those who were discharged home, p=0.013. A significantly higher proportion of study participants with secondary education or above were discharge home compared to those who died, 88 (35.2%) versus 16 (18.6%), p=0.012. The Mean modified Barthel index on admission was higher in participants who survived compared to those who died, 57.4 and 16.9 respectively, p< 0.001. Factors independently associated with mortality were total dependency on admission (modified Barthel index ≤20), adjusted OR 15.43, 95% CI: 7.523 – 31.664 (p<0.001) and male sex, adjusted OR 1.89, 95% CI: 1.012 – 3.528, p=0.046 (Table 4).

**Table 4:**
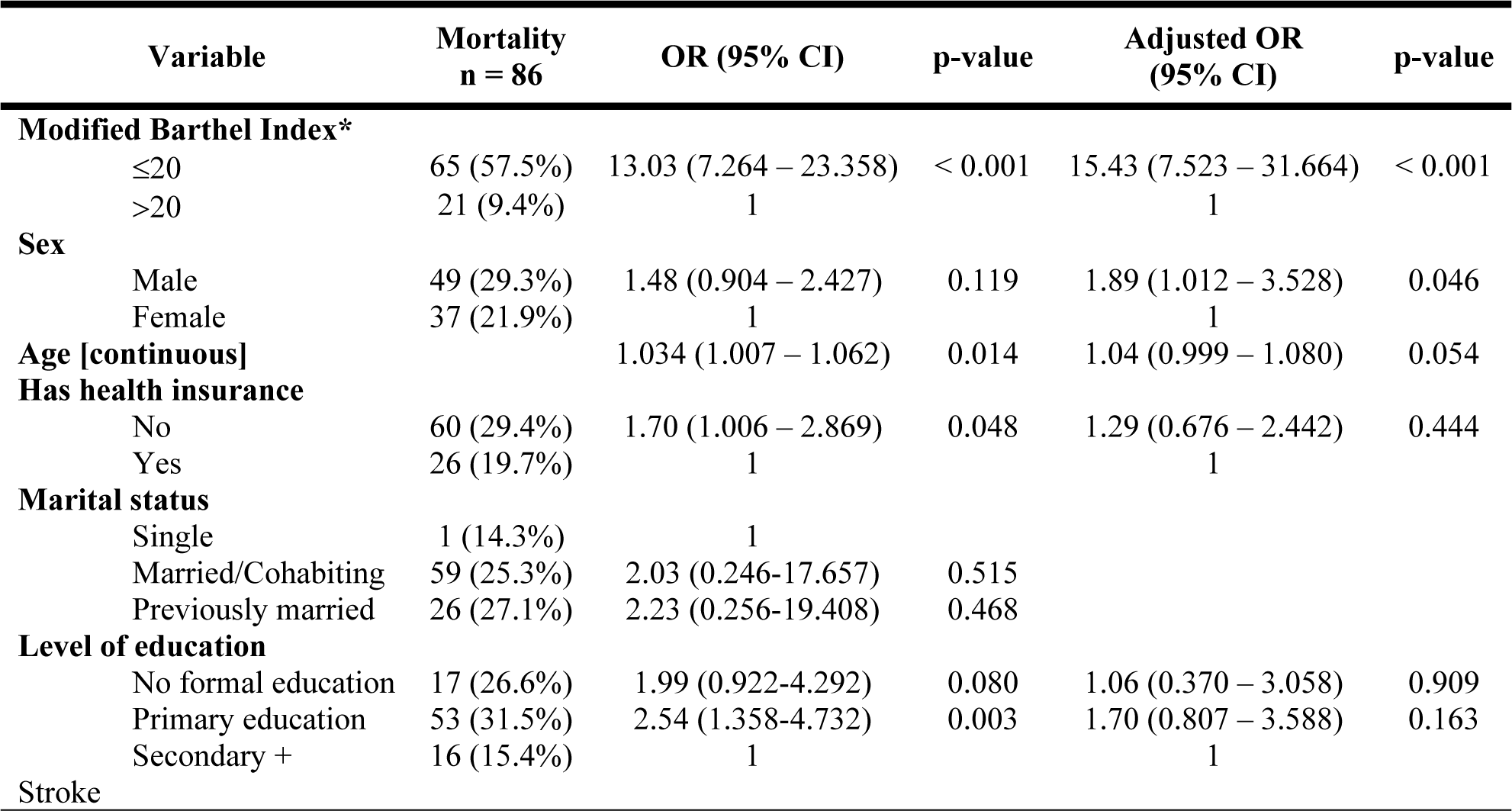
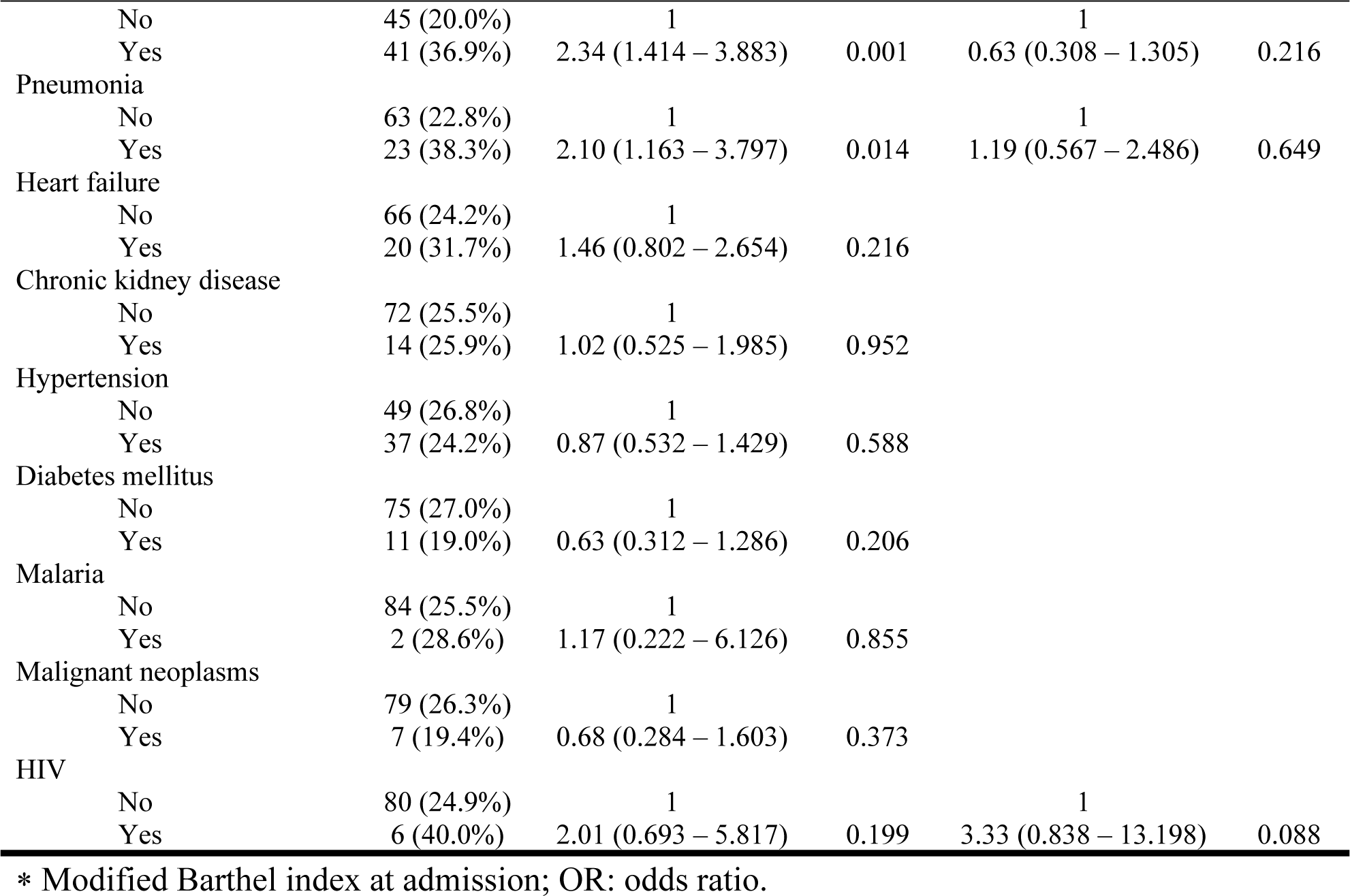
Factors associated with mortality among the 336 study participants.

## Discussion

Elderly patients comprised a third of all admissions through the emergency medicine department into JKCI and MNH medical wards despite constituting 8.3% of the individuals aged 15 years and above in the Tanzanian population [11]. The observed proportion of elderly admissions is double what was observed in the late 1990s in the same hospital [12]. The increased proportion of elderly medical admissions at MNH/JKCI may be because of several factors. Firstly, the number of elderly people in the general population has increased in the last three decades. This went hand in hand with increased life expectancy[1]. Secondly, elderly people are facing an increased burden of noncommunicable diseases (NCDs). Between 1990 and 2010 NCDs increased by 46 percent in sub-Saharan Africa [13]. The mean age of the study participants was similar to other recent studies among elderly patients in Africa [5]. In this study, there was a balance between males and females admitted contrary to previous studies which indicated males forming a greater proportion of hospital medical admissions [2–4,12]. All this has been occurring despite the fact that females tend to live longer than men and are more than men [14,15]. Findings from our study might reflect a change in social dynamics such as improved gender equity in access to health care. We have shown that literacy rate among the elderly was higher than that reported by previous studies. While studies done two to three decades ago had shown the majority of elderly patients to have no formal education[2,3], in this study 81% had at least primary education which may be the result of education policies that were implemented by the Tanzanian government to ensure primary education for all.

We found that the proportion of study participants with comorbidity is twice what was observed in the same setting three decades ago [2]. In addition, more than three-quarters of the diagnoses were noncommunicable diseases. Similar findings have been reported from other countries in sub-Saharan Africa [3,5,16]. In this study, the most frequent single diagnoses were hypertension, stroke and heart failure. These findings were expected given that aging is associated with increasing arterial stiffness and vascular resistance which leads to increasing blood pressure, strokes and myocardial ischemia, declining glomerular filtration rate and impaired response to antidiuretic hormone [17–19]. In addition, the presence of one of these diseases may be is a risk factor for another, for example, the presence of hypertension is a risk factor for heart failure or the presence of diabetes mellitus and hypertension form a significant risk for chronic kidney disease and stroke [20,21]. Indeed similar pattern of diseases among elderly medical admissions has been reported from other centers[3–5]. Infectious diseases like malaria formed a smaller proportion of admissions. The same trend was observed in previous studies [2,3,5] and may be due to the fact that malaria incidence is declining [22,23]. HIV diagnosis rate observed is consistent with the current HIV prevalence of 4.7% in Dar es Salaam adult population [24]. Lower HIV diagnosis rates ranging from 0 – 1.1% were reported in studies involving teaching hospitals in Nigeria, Sudan and northern Tanzania [5].

The median duration of hospitalization was similar to that reported from another tertiary center in Tanzania [5]. However, compared to studies conducted in other centers in Africa and elsewhere [4,5], the duration found in our study was shorter. This is reflected in the finding that 8.4% of the study participants discharged home had total dependency in ADL. Given the scarcity of home-based care services for diseases other than HIV/AIDS, elderly patients who are discharged home with such severe disability may be at increased risk to unfavorable outcomes including death. Since the duration of hospital stay is often a trade-off between the standard of care and socioeconomic burden on the patient and the health care system, a study is needed to evaluate and formulate optimum criteria for discharge. The fact that there is usually a supportive social fabric in Africa but not in Europe might also influence decisions on when to discharge patients from the hospital.

We found that more than 50% of deaths among elderly patients admitted to the medical wards occurred during the first 3 days of hospitalization. High initial mortality has also been reported in another study from Nigeria [4]. The high mortality, especially in the first few days of hospitalization, could be due to delays in seeking medical care and/or severity of the presenting clinical condition. Indeed, the in-hospital mortality was lower among patients with health insurance compared to those without (p=0.046). We believe that possessing a health insurance may be associated with better and early access to hospital care.

The observed in-hospital mortality of 25.6% is similar to that reported from other centers in Tanzania and other African countries [5,25]. However, the mortality was four times higher compared to in-hospital mortality in the United Kingdom [5]. The observed difference may be due to economic and other barriers to accessing health care leading to delays in diagnosis and treatment. In this study, over 60 % of the study participants had no health insurance and therefore were required to pay cash before they could access health care. This creates a major barrier to economically disadvantaged people. The high early mortality also signals the need for reviewing acute management of medical conditions in elderly patients.

The most important factor associated with mortality was total dependency on activities of daily living at admission similar to the findings of previous studies [26–28].

## Strength of the study

This study had an adequate sample size to provide current data on disease burden, diagnosis, and factors associated with the disease outcome in elderly patients admitted in the two hospitals. The prospective recruitment of the study participants ensured completeness of data obtained. Use of ICD-10 allowed for standardized coding and reporting of diseases. The Modified Barthel ADL index recommended for use in other settings has been used successfully in this study to evaluate functional status at admission and at discharge. Its prognostic significance has been shown.

## Study limitations

Data for this study including assessment of functional status using ADL scores was done in experimental/ non-routine conditions, in tertiary hospital facilities; feasibility of performing such assessment in routine conditions may be different.

The findings of this study are from tertiary level care facilities and may not be applicable to lower level care facilities. The study was not designed to follow-up individuals outside the hospital and hence doesn't provide outcomes beyond the date of hospital discharge to determine, for example, if the discharges are premature or not.

## Conclusions

We have shown that elderly patients constitute a significant proportion of the medical admissions at MNH and JKCI. While major communicable diseases such as malaria, HIV and tuberculosis constituted only a small proportion of the diagnoses; noncommunicable diseases accounted for over three-quarters of the diagnoses among elderly medical patients and were associated with high mortality especially in the first week of hospitalization. Modified Barthel index score ≤20 at admission was an independent risk factor for a poor in-hospital outcome and can, therefore, be used to identify patients who should be given special attention.

## Acknowledgement

The authors are grateful to Emanuel Pyuza for his assistance in participant recruitment and data collection.

## Author contribution

Conceptualization, B.T., K.P.; Methodology, B.T., K.P.; Formal analysis, B.T., K.P., P.M.; Investigation, B.T.; Writing – Original Draft, B.T., K.P., P.M.; Writing – Review & Editing, B.T., P.M., K.P.

## Supporting information captions

**S1 Table**. ICD-10 diagnosis categories: frequencies

